# An *in vivo* fitness gene of *Toxoplasma*, MIC11, is essential for PLP1-mediated egress from host cells

**DOI:** 10.1101/2025.03.24.644834

**Authors:** Yuta Tachibana, Miwa Sasai, Hidetaka Kosako, Eizo Takashima, Vern B Carruthers, Dominique Soldati-Favre, Masahiro Yamamoto

## Abstract

After invasion and replication, intracellular pathogens must egress from infected host cells*. Toxoplasma gondii* facilitates this process by permeabilizing host cells by releasing perforin-like protein 1 (PLP1) through induced microneme secretion. However, the precise mechanism of host cell permeabilization remains enigmatic. Here, we identified the secretory microneme protein MIC11 as a key factor for membrane disruption. A CRISPR-based *in vivo* screen revealed several genes including MIC11 as an essential gene for virulence. Deletion of MIC11 resulted in severe defects in both membrane rupture and egress. Scanning mutagenesis identified functional motifs in MIC11, and mechanistic analyses demonstrated that MIC11 directly associates with PLP1, regulating its activity in membrane disruption. The MIC11 paralogue MIC22 compensated for MIC11 deletion, suggesting a conserved mechanism of egress in the feline-restricted stages of *T. gondii*. The discovery of MIC11 advances the understanding of how parasites disrupt host cells to facilitate rapid egress and successful dissemination.

## Introduction

Intracellular bacteria and protozoa utilize various strategies to exit infected cells, a process known as egress^1^. Egress is a critical step for the obligate intracellular apicomplexan parasites, such as *Toxoplasma gondii* and *Plasmodium falciparum*, which cause human disease worldwide and are serious problems in public health^2^. This process relies on a dynamic interaction between host and pathogen factors and in such apicomplexans, it typically leads to the lysis of the host cell, resulting in detrimental effects for the host.

In *T. gondii*, egress is initiated by a signaling cascade that triggers microneme secretion causing the disruption of the vacuolar membrane surrounding parasites^3,4^. Recent studies have markedly advanced our understanding of the key regulators, such as guanylate cyclase (GC)^5^, protein kinase G (PKG)^6^, calcium-dependent protein kinases 1 and 3 (CDPK1 and CDPK3)^7–10^, and diacylglycerol kinase-1 (DGK1)^11^. While rhoptry secretion is not required for egress^12^, a dense granule phospholipase, lecithin-cholesterol acyltransferase (LCAT), has been shown to contribute to egress in *T. gondii*^13,14^. Beside responding to extrinsic signals, the parasite also produces its own signal via the dense granule protein diacylglycerol kinase 2 (DGK2) to control natural egress^15^. Micronemes are small secretory organelles localized at the apical end of the parasite^16^. They release a variety of cargo proteins known as microneme proteins (MICs), which include adhesins, proteases, and perforins essential for the parasite’s invasion, egress, and survival^17^. *T. gondii* is estimated to contain over 40 MICs^18^, but many of which are still functionally uncharacterized.

Perforin-like protein 1 (PLP1), a member of MICs, is widely conserved among the apicomplexans and is essential for egress in *T. gondii*^19^. Parasites lacking PLP1 exhibit significantly impaired egress from host cells and a complete loss of virulence in mice, highlighting the critical role of efficient egress in the *in vivo* fitness of *T. gondii*. Discharged PLP1 likely forms pores on the parasitophorous vacuole membrane (PVM), a membranous vacuole where parasites proliferate, leading to membrane permeabilization and disruption^20^. A current model supported by genetic, biochemical, and structural studies suggests that PLP1 alone is sufficient for membrane disruption^19,21–24^. Although a putative micronemal metalloprotease toxolysin 4 (TLN4) is implicated in egress, its contribution is limited^25^. Thus, PLP1 is currently thought to be the only effector protein that is both sufficient and necessary for membrane disruption, and the precise molecular mechanisms underlying this process are still poorly understood.

A genome-wide *in vitro* screen in human fibroblasts revealed many fitness-conferring genes during the lytic cycle^26^, but paradoxically found PLP1 to be dispensable. This discrepancy likely results from the screening protocol, which bypasses egress by mechanically lysing and passaging the host cells or delayed egress of ΔPLP1 parasites. In contrast, *in vivo* CRISPR screens, free from such interventions, consistently identified PLP1 as crucial for parasite fitness^27,28^, suggesting they are more effective for identifying egress factors. Such CRISPR/Cas9-based genetic screens *in vivo* have identified essential genes involved in various *T. gondii* processes^29,30^.

Here, we conducted a CRISPR-based *in vivo* screen targeting genes encoding proteins of the apical complex and the pellicle, leading to the identification of several previously unrecognized genes essential for *T. gondii* virulence. Among them, we identified microneme protein 11 (MIC11) as a key egress factor that directly interacts with PLP1 to mediate membrane disruption and parasite egress from host cells. Our findings challenge the long-standing model of *T. gondii* egress and provide a major conceptual advance in understanding the molecular mechanisms driving this critical process.

## Results

### *In vivo* CRISPR screen identifies MIC11 as the top-hit among the apical-pellicle library

The apical complex (including the conoid and apical secretory organelles) and the pellicle (inner membrane complex and parasite plasma membrane) are critical subcellular compartments of the apicomplexan parasites participating in motility, invasion, and egress^31,32^. This motivated us to search for genes essential for the parasite *in vivo* fitness by CRISPR-based screening of proteins localized to specific subcellular compartments, hypothesizing that novel egress factors might be among these fitness conferring genes.

We utilized a previously established *in vivo* CRISPR screen system using type I strain (RH) and inbred C57BL/6 mice^27^ **(Figure 1A)**. We designed a new guide RNA (gRNA) library targeting selected genes encoding the apical complex and the pellicle-localized proteins^18^ **(Figure 1B)**. RHΔhxgprt parasites were transfected with the gRNA library and were maintained in cell culture for four passages under the selection by pyrimethamine. The resulting mutant pool from the fourth passage was subsequently inoculated into the footpad of C57BL/6 mice at a dose of 10^7^ parasites per mouse. Seven days after infection, parasites were retrieved from the spleens and were inoculated to cell culture for one passage to expand parasites, yielding enough parasite genomic DNA for sequencing genome-integrated gRNAs. *In vitro* and *in vivo* fitness scores of each gene were calculated as the average log_2_ fold change of gRNA abundances between the input library and the fourth passage, or the fourth passage and the spleen, respectively. Our *in vitro* fitness scores were highly correlated with the genome-wide *in vitro* fitness scores in human foreskin fibroblasts (HFF)^26^ **(Figure S1A)**. High reproducibility of *in vivo* fitness scores was observed between independent infections in mice **(Figure S1B)**.

**Figure 1.**
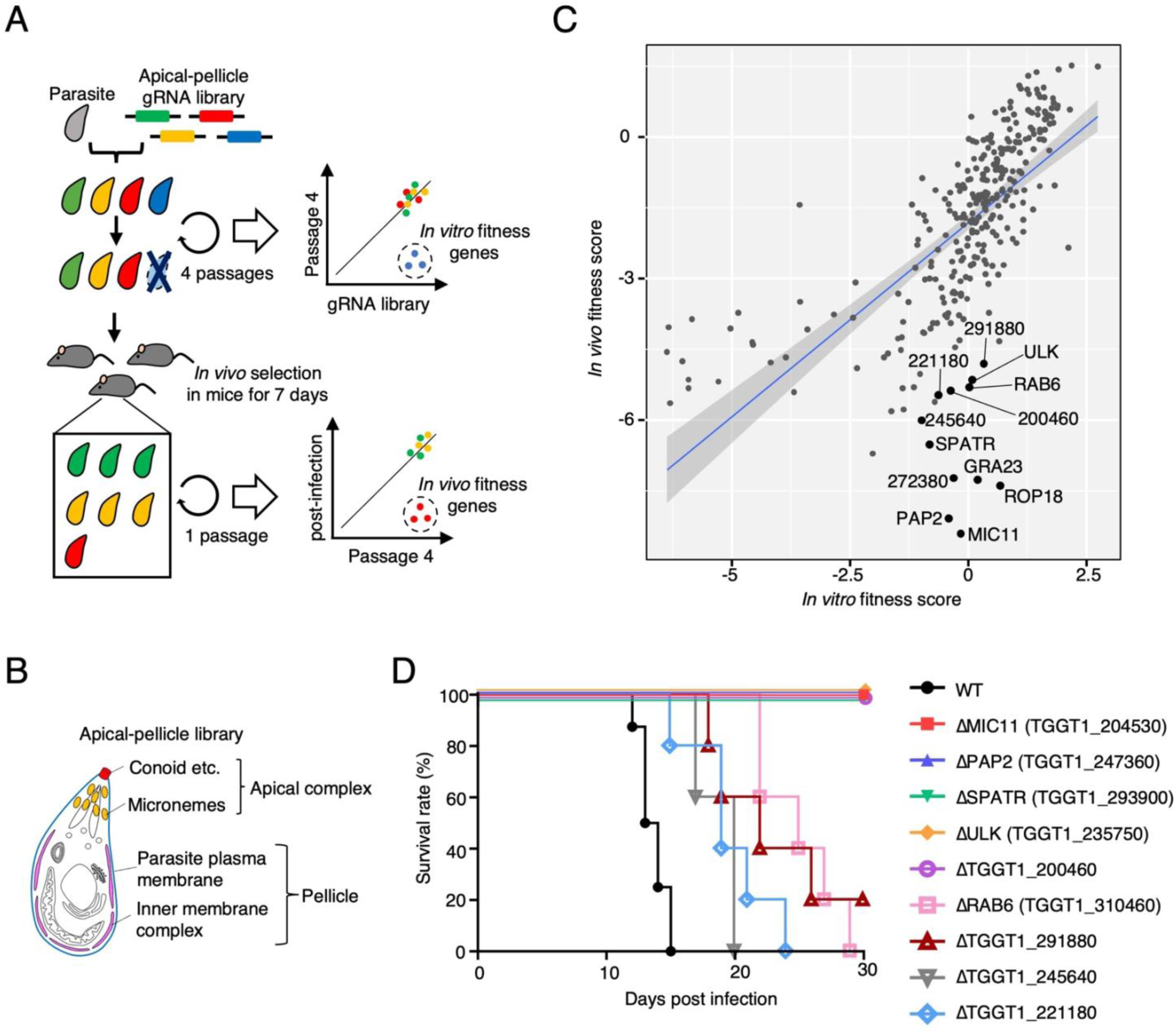
CRISPR screen identified novel genes required for virulence localized in the apical complex and the pellicle. (A) Schematic of *in vivo* CRISPR screens in mice. (B) Schematic of the apical-pellicle gRNA library. (C) A scatterplot of the screen results. Some top hits were labeled. ROP18 and GRA23 are control genes for *in vivo* fitness. (D) Virulence assays for candidate genes. 1000 parasites were injected into the footpad. WT (N = 8 mice) and others (N = 5 mice).

Comparing *in vitro* and *in vivo* fitness scores identified genes that contribute to parasite fitness during acute infection **(Figure 1C)**. The distance of each gene from the regression line was computed to identify top-ranking candidates **(Table S1)**. The control genes required for mouse infection, ROP18 and GRA23, were ranked highly. Some microneme-localized genes predicted by spatial proteomics were highly ranked, such as MIC11, SPATR, TGGT1_272380, and TGGT1_221180. MIC11 is also one of the top hits of another *in vivo* CRISPR screen^28^. SPATR was reported to be a component of the MIC complex essential for rhoptry discharge and contributes to virulence^33–35^. TGGT1_272380 was recently shown to be localized to the endoplasmic reticulum and dispensable for virulence^36^; thus, it was excluded from further analysis. Apical-localized gene products predicted by spatial proteomics, such as TGGT1_200460, TGGT1_291880, and TGGT1_245640, were also ranked highly. TGGT1_200460 was recently shown to be essential for virulence^28^. Pellicle-localized genes, such as PAP2 (TGGT1_247360), ULK kinase (TGGT1_235750), TGGT1_200460, and RAB6 (TGGT1_310460), were also ranked highly.

We selected nine genes displaying impaired *in vivo* fitness scores for validation, generating knockout parasites and inoculating them to the mouse footpad **(Figures 1D and S1C–S1D)**. Five knockouts (ΔMIC11, ΔPAP2, ΔSPATR, ΔULK, and ΔTGGT1_200460) were avirulent in mice. Four knockouts (ΔRAB6, ΔTGGT1_291880, ΔTGGT1_245640, and ΔTGGT1_221180) showed reduced virulence. Validation of these nine knockouts as important genes for virulence indicated our screen was reliable. In this study, we further describe the characterization of the top-hit MIC11 in the following sections.

### MIC11 is a microneme protein processed by aspartyl protease ASP3

MIC11 is a 16 kDa microneme protein conserved among coccidian parasites^37^, and its role remains unknown so far. MIC11 consists of a signal peptide, and two helical peptides, α-chain and β-chain, separated by an internal γ-propeptide. During maturation, the γ-propeptide is removed, and finally, the α-chain and β-chain are connected by a single disulfide bond, forming mature MIC11^37^ **(Figure S2A)**. We confirmed that MIC11 localized to micronemes and was secreted to the excretory-secretory antigens (ESAs) in a Ca^2+^-dependent manner **(Figures S2B and S2C)**.

Most secretory proteins of *T. gondii* are proteolytically cleaved either during their trafficking to secretory organelles or after exocytosis. *T. gondii* possesses several aspartyl proteases (ASPs), including ASP3, which was shown to process some rhoptry proteins and MICs, thus essential for parasite invasion and egress^38^. Previous proteomics data indicated that MIC11 was one of the potent candidates for ASP3 substrates^38^. Therefore, we used the anhydrotetracycline (ATc)-inducible ASP3 knockdown strain tagged with Ty epitope tag (ASP3-iKD) to assess if ASP3 serves as maturase for MIC11. Western blot analysis confirmed that ATc treatment for 48 h depleted ASP3 protein expression and as expected, its substrate MIC6 accumulated as an immature form. MIC11 showed a reduced amount of mature form **(Figure S2D)**, indicating ASP3-dependent processing at two putative sites in accordance with the previous study^38^ **(Figure S2A)**. Taken together, these results indicate that MIC11 is a soluble microneme protein processed by ASP3.

### MIC11 is essential for parasite egress and vacuolar permeabilization

Plaque assays were performed to assess the impact of MIC11 deletion on the lytic cycle and ΔMIC11 produced smaller plaques compared to WT parasites. The complemented ΔMIC11::MIC11-Spot-HA parasites, C-terminally tagged with Spot and HA epitope tags, restored normal plaque formation **(Figures 2A and S3A–S3B)**, confirming that MIC11 is required for the lytic cycle of *T. gondii*. Further investigation revealed that MIC11 deletion did not affect invasion or intracellular proliferation **(Figures S3C–S3D)**. Contrastingly, abnormal spherical structures where multiple parasites were entrapped, reminiscent of those observed in the ΔPLP1 parasite-infected cells^19^. Those spheres were not observed in WT or the complemented strains **(Figure 2B)**. Considering that PLP1 is necessary for egress, we performed induced egress assays following stimulation with the Ca^2+^ ionophore ionomycin to assess the role of MIC11 in egress. Most WT parasites egressed from the infected cells within 5 min, while ΔMIC11 parasites remained inside vacuoles indicating that induced egress was severely impaired in ΔMIC11 parasites. ΔMIC11::MIC11-Spot-HA parasites recovered the egress phenotype **(Figures 2C and 2D)**. Another stimulation, such as low pH, also triggers parasite egress from host cells^21,39^. We also performed low pH-induced egress assays and ΔMIC11 parasites exhibited a severe defect **(Figure S3E)**. After intraperitoneal injection, ΔMIC11 parasites lost virulence in mice, which was like ΔPLP1 parasites^19^, and ΔMIC11::MIC11-Spot-HA parasites restored virulence **(Figure 2E)**.

**Figure 2.**
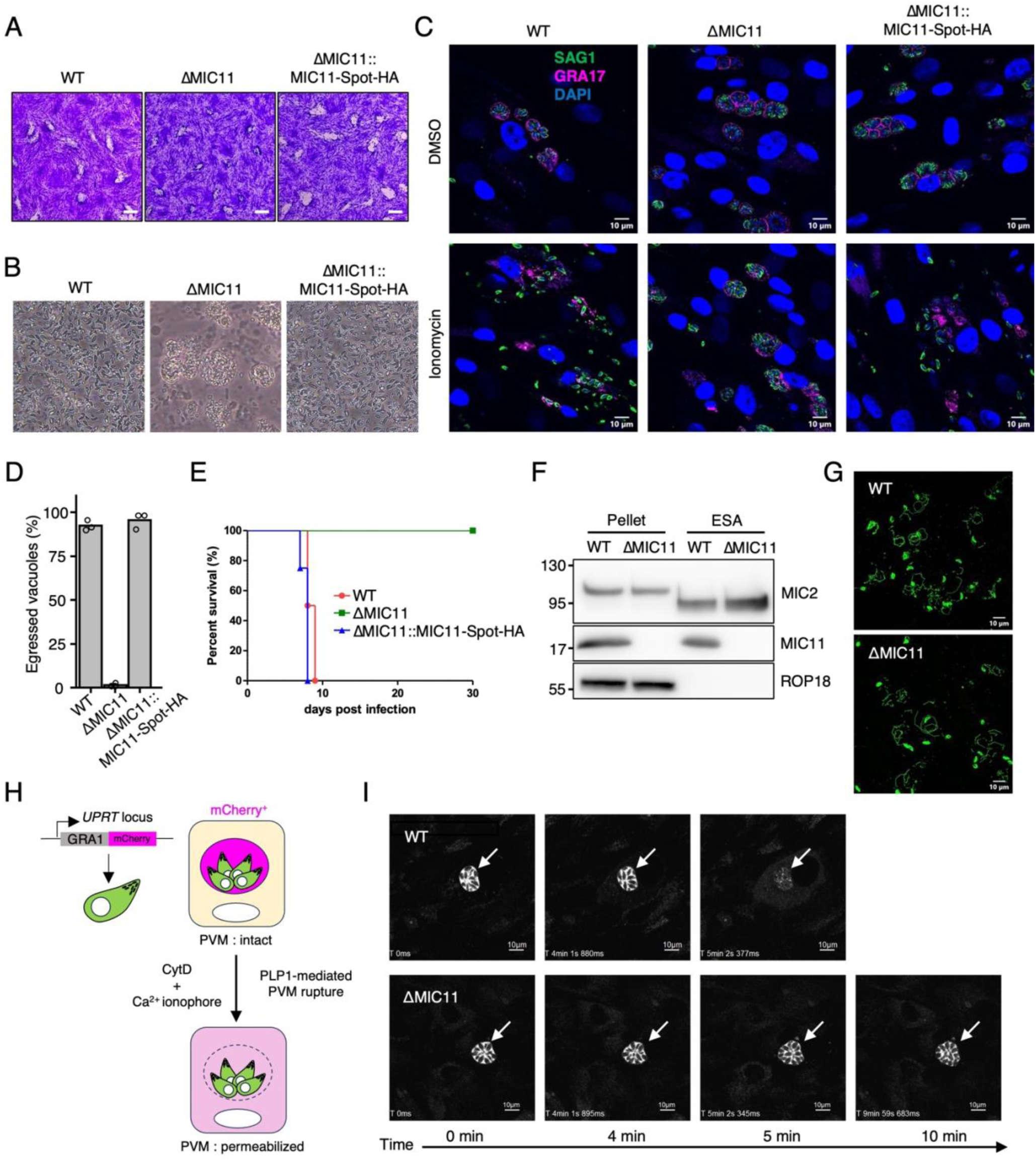
MIC11 is essential for parasite egress and PVM permeabilization. (A) Representative images of plaque assay of indicated strains. Scale bar = 1 mm. (B) Parasite cultures 72h post-infection. (C) Immunofluorescence images of ionophore-induced egress assay. SAG1 (green) and GRA17 (magenta) indicate parasites and PVs, respectively. (D) Ionophore-induced egress assay of indicated strains. Data are displayed as mean values (n = 3). (E) Virulence assays of indicated strains. 1000 parasites were injected intraperitoneally. N = 5 mice per group. (F) Microneme secretion assay. (G) Gliding motility assay. (H) Schematic of PVM permeabilization assay. (I) Time-lapse images of ΔMIC11 and WT parasites treated with cytochalasin D and ionomycin.

To unravel the mode of action of MIC11, we conducted deeper investigations to examine which steps are affected by MIC11 deficiency. Microneme secretion and gliding motility of ΔMIC11 parasites were normal **(Figures 2F and 2G)**. Transgenic parasites expressing GRA1-mCherry were generated to evaluate the PVM integrity. The PVM is intact in unstimulated conditions, and GRA1-mCherry resides in the PV. Parasites are then immobilized with the actin polymerization inhibitor cytochalasin D (CytD). Ca^2+^ ionophore treatment induces microneme secretion, and the discharge of PLP1 causes PV rupture. Finally, GRA1-mCherry diffuses from the PV to the host cytosol^19^ **(Figure 2H)**. Then, we monitored the permeabilization of the PVM after stimulation with ionomycin by using time-lapse imaging. WT parasites caused PV rupture after ionophore stimulation within 5 min. In contrast, no PV rupture was observed in ΔMIC11 vacuoles within 10 min **(Figure 2I)**. These findings demonstrate that defects in membrane permeabilization account for impaired egress in ΔMIC11 parasites.

### Scanning mutagenesis revealed functional motifs in MIC11

Next, we attempted to identify which regions of MIC11 are required for membrane permeabilization. MIC11 lacks homology to known proteins and has no apparent conserved domains other than a signal peptide. To unbiasedly assess the determinants for MIC11 localization and function, we performed scanning mutagenesis by replacing each set of five amino acid residues with five consecutive alanine^40^. We also generated point mutations C90A and C194A because a disulfide bond is formed between C90 and C194^37^. Mutant parasites were generated by transfecting MIC11 mutant vectors into ΔMIC11 parasites, yielding MIC11 mut_01–mut_35, C90A, and C194A, as we did previously for wild-type MIC11. Expression and localization of MIC11 mutants were validated by immunofluorescence assays (IFA) **(Figure S4)**. Ionophore-induced egress assays were performed to assess the phenotype of each mutant parasite. The results mapped the critical and dispensable regions in MIC11 across the α-chain, β-chain, and γ-propeptide **(Figure 3A)**. The α-chain possesses immutable regions in the anterior and the middle, whereas the immutable regions are spreading throughout the β-chain. As expected, C90A and C194A mutations failed to restore the egress phenotype.

**Figure 3.**
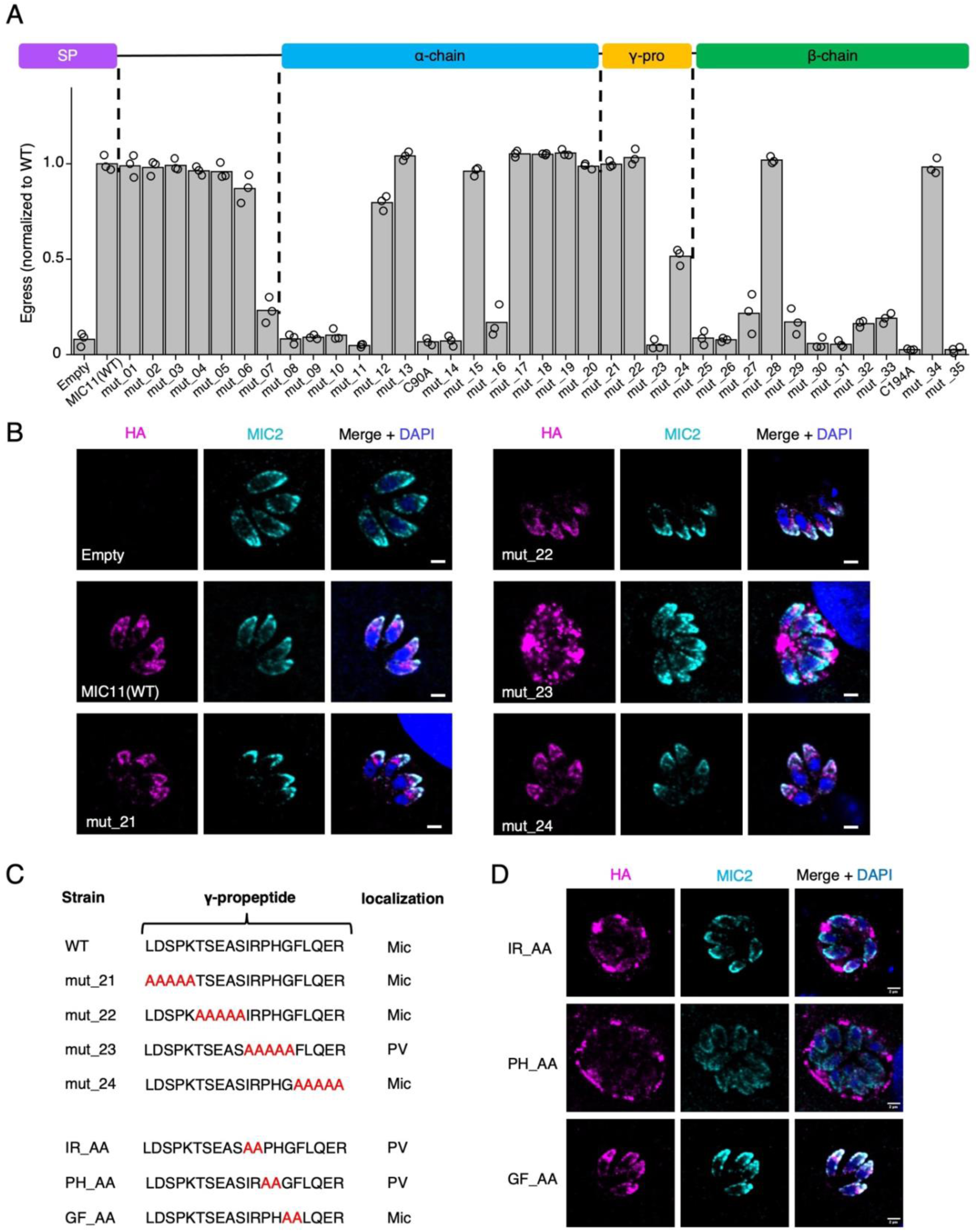
Scanning mutagenesis revealed that the gamma-propeptide of MIC11 is essential for microneme localization. (A) Top, Schematic of MIC11 domain. Bottom, induced egress of wild-type and 5-Ala substituted mutants of MIC11. Each sample percentage of egressed vacuoles was normalized against wild-type MIC11 control. (B) MIC11 localization in indicated strains. MIC11(mut_23) is not colocalized with MIC2. (C) The γ-propeptide sequences and MIC11 localization of wild-type and indicated mutants. (D) MIC11 localization in indicated strains.

We found that the mut_23 (IRPHG^137–141^ to AAAAA) showed abnormal localization of MIC11 among the γ-propeptide mutants, seeming to be localized in the PV, not in micronemes **(Figures 3B and 3C)**. We hypothesized that there is a determinant for trafficking to the microneme within the γ-propeptide. Dissection with two amino residue substitutions **(Figure 3C)** showed abnormal MIC11 localization with the IR_AA and PH_AA mutants, while the GF_AA mutant showed normal microneme localization **(Figure 3D)**. These results demonstrated that the IRPH sequence in the γ-propeptide is essential for trafficking to micronemes.

### MIC11 directly interacts with PLP1 to facilitate parasite egress

To understand how MIC11 contributes to the PVM disruption, we performed co-immunoprecipitation mass spectrometry (IP-MS) analysis. Comparing the enrichment in protein abundance between immunoprecipitations from ΔMIC11::MIC11-Spot-HA parasites over the parental ΔMIC11 parasites revealed several MICs: PLP1, MIC5, MIC10, MIC20, and MIC21 **(Figure 4A and Table S2)**. It is widely known that many MICs interact with other MICs to form a functional complex^17^. Except for PLP1, the other four MICs have not been reported to be associated with parasite egress^41–44^. Our *in vivo* CRISPR screen results indicated that PLP1 is essential for parasite fitness, while the other four MICs were not essential, at least by their single deletion^27^ **(Figure 4B)**. Hence, we generated ΔPLP1 parasites and MICs quadruple knockout parasites (ΔMIC5ΔMIC10ΔMIC20ΔMIC21, referred to as QKO) **(Figures S5A–S5C)**. As reported previously^19^, ΔPLP1 parasites showed severely impaired egress and virulence, while QKO parasites showed normal egress and virulence comparable to WT parasites **(Figures 4C and 4D)**. These results suggest that the functionally most relevant interaction is between MIC11 and PLP1. To investigate the relationship between MIC11 and PLP1 further, double knockout parasites for MIC11 and PLP1 (ΔMIC11ΔPLP1, referred to as DKO) were generated. We complemented DKO parasites with either MIC11-HA, PLP1-Ty, or both **(Figures 4E and 4F)**. Only the double-complemented DKO::MIC11-HA::PLP1-Ty parasites restored the egress phenotype **(Figure 4G)**, indicating that both MIC11 and PLP1 are necessary for parasite egress.

**Figure 4.**
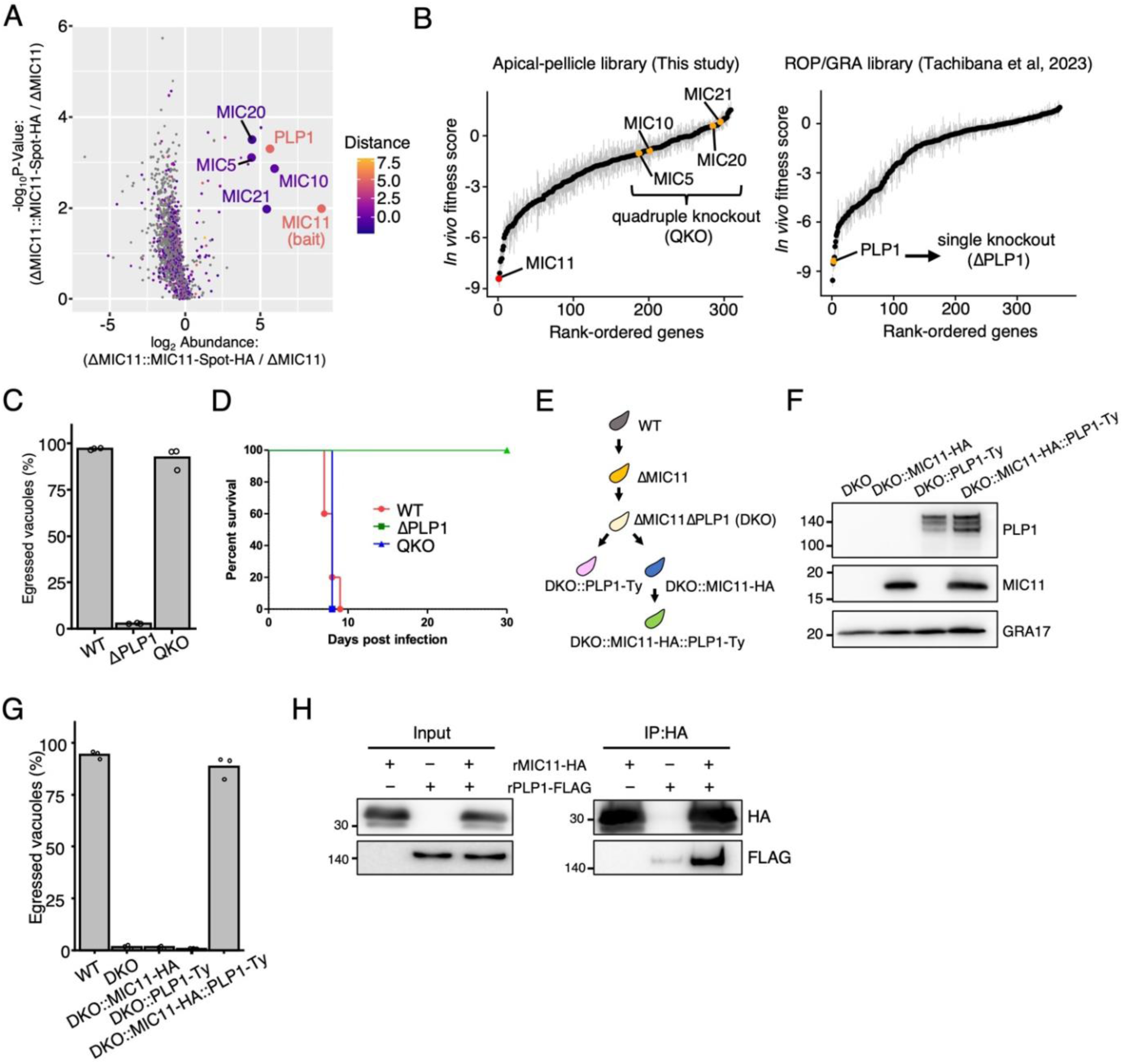
MIC11 interacts with another microneme protein PLP1 facilitating parasite egress. (A) Several microneme proteins (MIC5, MIC10, MIC20, MIC21 and PLP1) were co-immunoprecipitated with MIC11-Spot-HA. The color scale indicates the distance for each gene in *in vivo* CRISPR screens. (B) Ranking plots of current and previous *in vivo* CRISPR screens indicate *in vivo* essentialities of each MIC. (C) Ionophore-induced egress assays of WT, ΔPLP1, and QKO parasites. Data are displayed as mean values (n = 3). (D) Virulence assay of indicated strains. 1000 parasites were injected intraperitoneally. N = 5 mice per group. (E) Schematic representation of double knockout and complementation of MIC11 and PLP1. (F) Western blot showing successful complementation of MIC11 or PLP1. (G) Ionophore-induced egress assays of indicated strains. Data are displayed as mean values (n = 3). (H) Co-immunoprecipitation of recombinant MIC11 and PLP1.

To examine whether MIC11 and PLP1 interact directly or indirectly, we used the wheat germ cell-free protein synthesis system to obtain epitope-tagged recombinant proteins, rMIC11-HA and rPLP1-FLAG. Analysis of the rMIC11–rPLP1 binding by co-immunoprecipitation demonstrated that both proteins were co-precipitated **(Figure 4H)**. Collectively, MIC11 and PLP1 likely interact directly and form a complex, facilitating parasite egress.

### MIC11 is required for the function of discharged PLP1

We further investigated the impact of MIC11 on PLP1 trafficking/processing/secretion and vice versa. We found that MIC11 localization is not affected by PLP1 deletion **(Figure 5A)** and MIC11 deletion did not affect PLP1 localization either **(Figure S6A)**. To assess the secretion and processing of PLP1, we tagged PLP1 with a myc-epitope tag at the C-terminus since PLP1 is N-terminally processed after exocytosis^22^ **(Figure S6B)**. PLP1 secretion and processing appear to be unaffected by MIC11 deletion **(Figure 5B)**. These results suggested that ΔMIC11 parasites could not permeabilize the membrane despite normal secretion of PLP1 and hence hypothesized that MIC11 regulates the function of PLP1 after exocytosis to disrupt the membrane.

**Figure 5.**
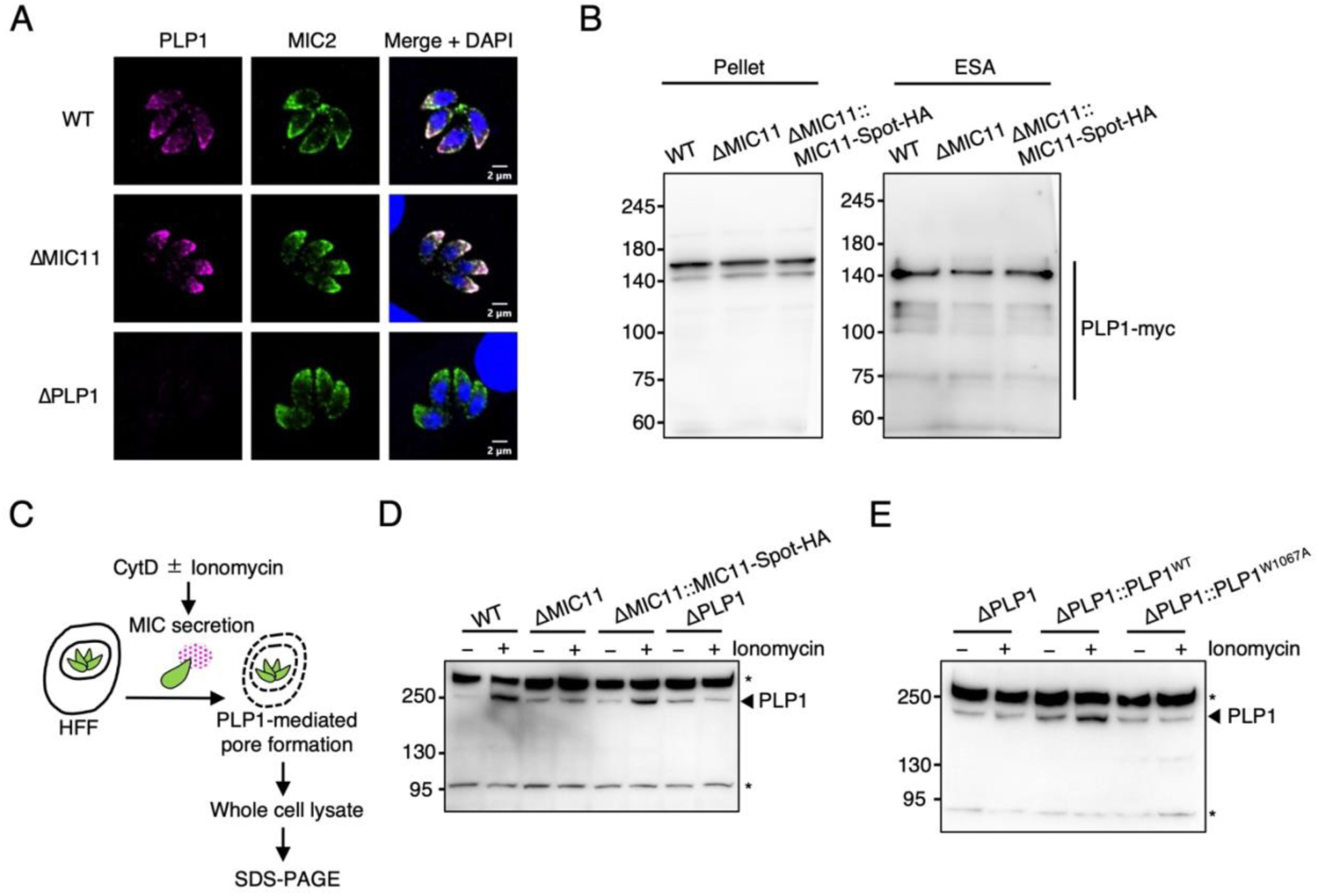
MIC11 is involved in PLP1 activity post-exocytosis. (A) Immunofluorescence images stained with anti-PLP1 (magenta) and anti-MIC2 (green) antibodies. (B) Immunoblots showing PLP1 secretion and processing of indicated strains. (C) Schematic representation showing detection of PLP1 dynamics induced by ionophore treatment. (D) Western blot showing the higher size of PLP1 in ionophore-treated WT and ΔMIC11::MIC11-Spot-HA strains. The arrow indicates PLP1 signal and there is a non-specific band of the same size. Asterisks indicate non-specific bands. (E) Western blot showed a higher size of PLP1 in ionophore-treated wild-type PLP1-expressing parasites but not in the loss of function mutant of PLP1. The arrow indicates PLP1 signal and there is a non-specific band of the same size. Asterisks indicate non-specific bands.

To capture the dynamics of discharged PLP1 during membrane permeabilization, we treated parasite-infected host cells with CytD to immobilize parasites and stimulated them with ionomycin. The whole cell lysate, including host cells and parasites, was subjected to western blotting for PLP1 detection **(Figure 5C)**. Notably, we observed a higher size of PLP1 band around 250 kDa in ionophore-stimulated WT parasite-infected cells. This band is not present in unstimulated WT parasite-infected cells, stimulated ΔMIC11-infected host cells, or stimulated ΔPLP1-infected host cells but is present in stimulated ΔMIC11::MIC11-Spot-HA-infected host cells **(Figure 5D)**, suggesting a close correlation with egress.

We further attempted to characterize the higher-size band of PLP1. Previous studies have shown that PLP1 is inserted into a lipid bilayer via a protruding hydrophobic loop with an exposed tryptophan residue at its tip, and the tryptophan mutation (W1067A) loses its ability to bind to membranes and permeabilize^23,24^. Consistent with a previous study^24^, we confirmed that ΔPLP1 parasites complemented with mutant PLP1 (ΔPLP1::PLP1_W1067A_) failed to egress in response to ionomycin **(Figure S6C)**. Further, we did not observe the higher-size PLP1 band in ΔPLP1::PLP1_W1067A_-infected cells **(Figure 5E)**, indicating that the higher-size PLP1 band is correlated with membrane binding and subsequent pore-formation mediated by PLP1. These results demonstrated that MIC11 is critical to the PLP1-mediated process during membrane permeabilization.

### A putative MIC11 paralogue is expressed in the feline-restricted stages of *T. gondii*

Sexual reproduction of *T. gondii* only occurs during the infection of the definitive feline host^45^. The parasites of feline-restricted stages need to egress from the intestinal cells, as do tachyzoites in the intermediate hosts. However, PLP1 and MIC11 are not expressed during the feline-restricted stages^46^. A previously proposed model suggested that PLP2, a PLP1 paralogue, is involved in egress in the feline gut^47^. This motivated us to search for a homologue of MIC11 expressing during the feline-restricted stages. We found one gene, TGME49_295662, encoding an uncharacterized hypothetical protein that shares 25% amino acid sequence similarity with MIC11 **(Figure 6A)**. TGME49_295662 possesses a signal peptide, α-chain, β-chain, and γ-propeptide-like sequences. Within the γ-propeptide-like region, an IAPH sequence resembles the IRPH microneme trafficking determinant in MIC11 **(Figure 6B)**. In contrast to MIC11, TGME49_295662 is highly expressed in merozoites^46^ **(Figure 6C)**.

**Figure 6.**
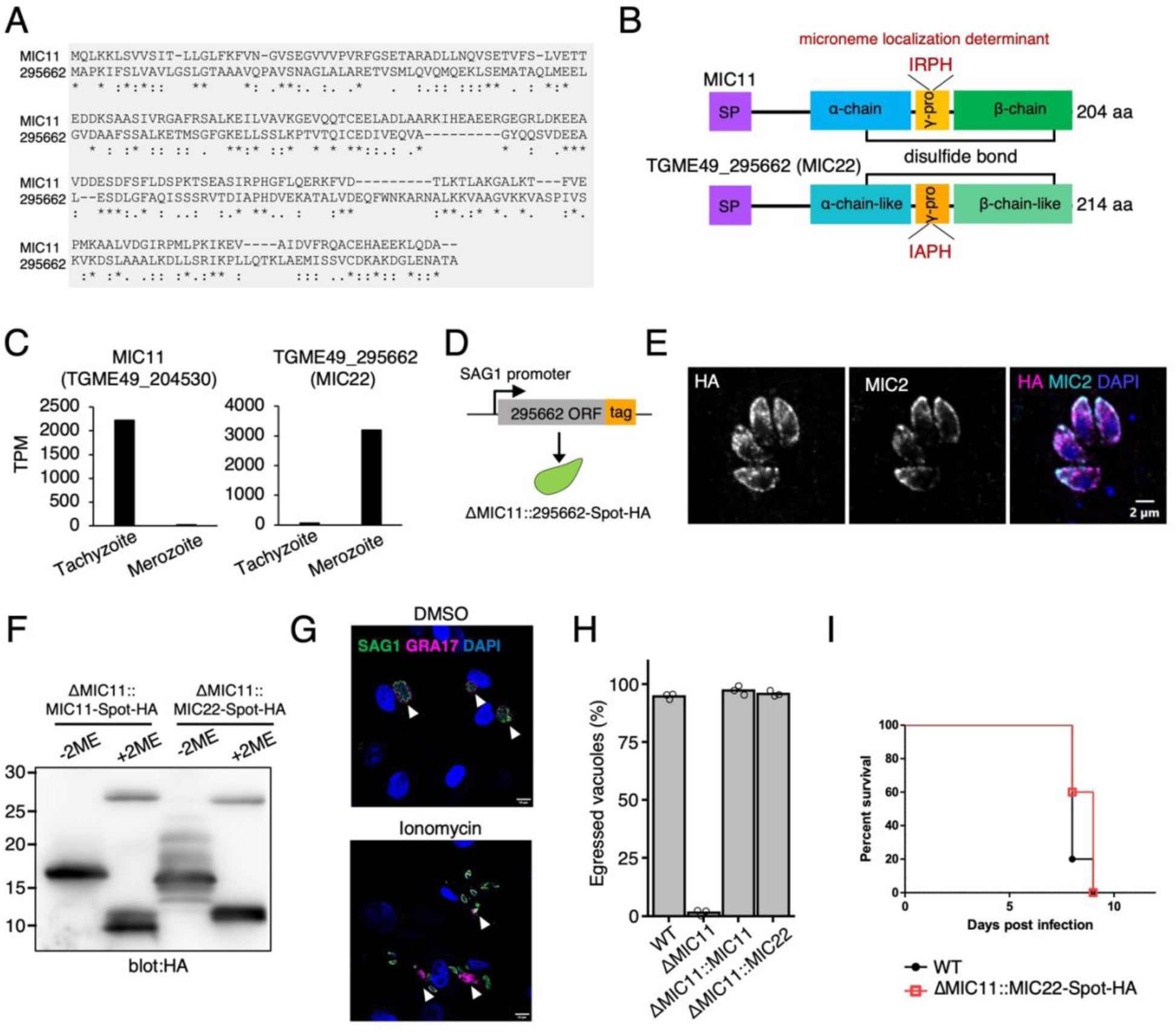
A merozoite-specific paralog of MIC11, MIC22, compensates MIC11 deletion. (A) Sequential alignment of MIC11 and TGME49_295662 (MIC22) by MAFFT. (B) Schematic of MIC11 and TGME49_295662 (MIC22). (C) RNA-sequence data (Hehl et al., 2015) of MIC11 and TGME49_295662 mRNA expression in tachyzoite and merozoite stages. (D) Schematic of transgenic parasites expressing MIC22 driven by SAG1 promoter in tachyzoite. (E) Immunofluorescence images of ΔMIC11::TGGT1_295662-Spot-HA showing microneme localization of TGGT1_295662-Spot-HA (magenta) colocalizing with MIC2 (cyan). (F) Immunoblots of parasites expressing MIC11-Spot-HA or MIC22-Spot-HA under non-reducing (−2ME) and reducing (+2ME) conditions. (G) Representative images of ionophore-induced egress assay of ΔMIC11::MIC22-Spot-HA parasites. SAG1 (green) and GRA17 (magenta) stain parasites and PVM, respectively. Arrowheads indicate occupied or egressed vacuoles. (H) Ionophore-induced egress assay of indicated strains. Data are displayed as mean values (n = 3). (I) Virulence assay of indicated strains. 1000 parasites were injected intraperitoneally. N = 5 mice per group.

To characterize the putative paralogue of MIC11 in tachyzoites, the open reading frame (ORF) of TGGT1_295662 was tagged with Spot and HA epitopes and expressed under the control of the SAG1 promoter in ΔMIC11 parasites **(Figure 6D)**. IFA revealed that TGGT1_295662-Spot-HA localized to micronemes in tachyzoites **(Figure 6E)**. ΔMIC11::MIC11-Spot-HA and ΔMIC11::295662-Spot-HA were subjected to western blotting under non-reduced or reduced conditions. The band of TGGT1_295662-Spot-HA showed a pattern like MIC11-Spot-HA **(Figure 6F)**, suggesting that a mature form of TGGT1_295662 consists of α-chain and β-chain, connected by a disulfide bond between C93 and C202 **(Figure 6B)**. Thus, we named TGGT1_295662 as microneme protein 22 (MIC22). We examined whether MIC22 compensates for MIC11 deficiency since MIC11 and MIC22 showed similar properties. The ΔMIC11::MIC22-Spot-HA strain fully restored the egress and virulence completely **(Figures 6G–6I)**. These results strongly suggest that PLP1-MIC11-mediated egress in tachyzoite is carried out by PLP2-MIC22 in feline-restricted stages.

## Discussion

Exit from the host cells is a critical process for intracellular pathogens, known as egress. In this study, we performed an *in vivo* CRISPR screen targeting genes encoding proteins of the apical complex and the pellicle to identify novel egress factors beyond PLP1. This led to the identification of MIC11, a secretory protein uncharacterized over the past 20 years, as a fundamental factor for egress in *T. gondii*. A previous study showed that PLP1 is sufficient for membrane disruption based on the lytic activity of recombinant PLP1 protein^22^. Our results support a new model wherein PLP1 alone is insufficient *in cellulo* and requires MIC11 to mediate membrane disruption.

Researchers have developed various genetic screening approaches to identify critical factors involved in *T. gondii* egress, leading to the discovery of multiple egress-associated genes^48–53^. However, despite its well-established role in egress, PLP1 has been missed in some screenings. We and others have conducted *in vivo* CRISPR screens and successfully identified MIC11 and PLP1 as *in vivo* fitness-conferring genes^27,28^. This suggests that the *in vivo* CRISPR screen serves as the phenotypic screen for egress-related genes, and additional yet unidentified egress factors may be present among the screening hits. Furthermore, host-derived factors such as calpain and G protein-coupled receptors have been implicated in the egress of apicomplexan parasites^54,55^. Understanding the complex host-parasite interactions that regulate egress remains a key focus for future investigations.

The RH strain tachyzoites naturally egress around 48 h post-infection in cell culture^15^. However, it was reported that *T. gondii* may egress earlier from infected host cells *in vivo*^56^. Additionally, external environments, such as cytokine stimulation or inflammation, can trigger premature egress of *T. gondii*^3^. Since ΔMIC11 or ΔPLP1 parasites exhibit a complete loss of virulence in mice, efficient egress appears to be essential for parasite fitness *in vivo*. The precise role of efficient egress in *in vivo* settings remains an important topic for future investigation.

Compared to tachyzoites and bradyzoites in intermediate hosts, feline-restricted stages are less explored. Interestingly, many secretory proteins show dynamic changes in the expression between tachyzoites and merozoites^46^. Our findings reveal functional redundancy between MIC11 and MIC22. While the role of PLP2 in feline-restricted stages remains unknown, MIC22 and PLP2 may function as key mediators of egress in merozoites. Recently developed methods of *in vitro* production of merozoite-like *T. gondii* would enable further exploration^57,58^.

In *Plasmodium* spp., proteases and phospholipases play more critical roles than perforins in egress during the asexual blood stages^59,60^. However, the *Plasmodium* genome encodes five PLP genes (PPLP1–5)^61^. Unlike *T. gondii*, *Plasmodium* parasites do not appear to possess MIC11-like genes. Future studies should investigate whether PPLPs alone are sufficient for membrane disruption or if additional factors contribute to this process. How the PVM is ruptured by apicomplexan parasites is a key question in understanding the molecular basis of parasite egress.

A limitation of this study is that it is still unclear how MIC11 contributes to membrane disruption in concert with PLP1. Membrane pore formation mediated by pore-forming proteins typically involves multiple steps, including membrane binding, insertion, oligomerization, and pore-formation^62,63^. Future studies with structural insights will be essential to elucidate the precise contribution of MIC11 during this process.

In summary, this study shows that MIC11 is the second factor indispensable for egress in *T. gondii* through direct interaction with PLP1. The identification of MIC11 paves the way for future studies on parasite egress and facilitates the development of promising therapeutic strategies aimed at preventing parasite egress and dissemination.

## Materials and methods

### Toxoplasma strains

All *T. gondii* strains were maintained in Vero cells by serial passage using RPMI (Nacalai Tesque) supplemented with 2% heat-inactivated fetal bovine serum (FBS; JRH Bioscience), 100 U/ml penicillin and 0.1 mg/ml streptomycin (Nacalai Tesque) in incubators at 37℃ and 5% CO_2_. The parental strains used in this study were RHΔhxgprt^27^ and RHΔhxgprtΔku80^64^. ASP3-iKD was described previously^38^.

### Host cell culture

Vero cells were maintained in RPMI (Nacalai Tesque) supplemented with 10% heat-inactivated FBS, 100 U/ml penicillin and 0.1 mg/ml streptomycin (Nacalai Tesque) in incubators at 37℃ and 5% CO_2_. HFFs were maintained in Dulbecco’s modified Eagle’s medium (DMEM) (Nacalai Tesque) supplemented with 10% heat-inactivated FBS, 100 U/ml penicillin and 0.1 mg/ml streptomycin (Nacalai Tesque) in incubators at 37℃ and 5% CO_2_.

### Mice

C57BL**/**6NCrSlc (C57BL/6N) mice were purchased from SLC. All animal experiments were conducted with the approval of the Animal Research Committee of Research Institute for Microbial Diseases in Osaka University.

### Antibodies

The following primary antibodies were used in the immunofluorescence assay, immunoblotting, or immunoprecipitation. Mouse anti-HA (MBL, M180-7), mouse anti-HA (MBL, 561-8), mouse anti-HA.11 (BioLegend, 901514), mouse anti-Ty (BB2), mouse anti-FLAG (Sigma, F3165), rabbit anti-myc (MBL, 562), mouse anti-GRA1 (TG17-43), mouse anti-MIC2 (T34A11)^65^, rabbit anti-MIC2, rabbit anti-MIC11, rabbit anti-MIC6^66^, rabbit anti-PLP1 (N-terminus)^22^, rabbit anti-PLP1, rabbit anti-GRA17^27^, rabbit anti-ROP18^67^, mouse anti-SAG1^68^, rabbit anti-GAP45^68^. We raised custom antibodies against synthetic peptides in rabbits as follows. MIC11 (RGEGRLDKEEAVDDESD), PLP1 (PTSKTMSSLKLAPVK), MIC2 (TKVVMEEEKETLV). The epitope identification, peptide synthesis, rabbit immunization, and serum collection were conducted by Cosmo Bio.

### Parasite transfection

Parental parasites were filtered and resuspended in transfection solution containing 2 mM ATP, 5 mM glutathione, DNA, and cytomix (10mM KPO_4_, 120mM KCl, 0.15 mM CaCl_2_, 5mM MgCl_2_, 25 mM HEPES, 2 mM EDTA). Parasites were electroporated in a 2-mm cuvette at 1500 V and 25 µF by GENE PULSER II (Bio-Rad). Stable transgenic parasites were selected with the appropriate antibiotics or sorted by cell sorter, cloned in 96-well plates by limiting dilution, and verified by genotyping, mRNA expression, or protein expression. The primers used for generating transgenic parasites are shown in **Table S3**.

### Generation of parasite transgenic lines

For generating knockout parasites, the specific gRNAs targeting each gene’s 5′ and 3′ ends were cloned into the pU6-Universal vector (Addgene #52694). The floxed HXGPRT construct was PCR amplified with 60 bp homology arms to each gene’s 5′ and 3′ UTRs. Parasites were transfected with two gRNAs and the PCR-amplified HXGPRT repair template. Transfected parasites were selected by 25 µg/ml mycophenolic acid (Sigma) and 50 µg/ml xanthine (Wako).

For generating quadruple knockout parasites (ΔMIC5ΔMIC10ΔMIC20ΔMIC21, QKO), parasites were transfected with eight gRNAs and the PCR-amplified HXGPRT, DHFR-TS, Venus, and mCherry repair templates targeting each gene. Transfected parasites were selected by 25 µg/ml mycophenolic acid, 50 µg/ml xanthine, and 3 µM pyrimethamine for serial passage. Stable resistant parasites were sorted for Venus and mCherry double positive parasites by BD FACS Aria III.

To generate endogenously tagged PLP1 strains, the myc-HXGPRT cassette was PCR amplified with 60 bp homology regions to the C-terminus of PLP1. The myc-HXGPRT repair template was co-transfected with a gRNA vector targeting the C-terminus of PLP1. Transfected parasites were selected by 25 µg/ml mycophenolic acid and 50 µg/ml xanthine.

To generate GRA1-mCherry expressing parasites, pUPRT-pTub-GRA1-mCherry plasmid was co-transfected with a gRNA vector targeting UPRT locus. Transfected parasites were selected by 10 µM fluorodeoxyuridine (Wako).

To complement PLP1 knockout parasites, PLP1 cDNAs (WT and W1067A) were amplified from the cDNA of the RH strain and cloned into the pUPRT vector. The pUPRT-PLP1-Ty vectors were co-transfected with a gRNA vector targeting UPRT locus. Transfected parasites were selected by 10 µM fluorodeoxyuridine.

To complement MIC11 knockout parasites, wild-type MIC11 cDNA was amplified from the cDNA of the RH strain. Mutated MIC11 cDNAs were synthesized by FASMAC. MIC22 ORF was amplified from the genomic DNA of the RH strain. Each cDNA was cloned into pHXGPRT vector.

ΔMIC11Δhxgprt parasites were transfected with the pHXGPRT-pSAG1-MIC11/MIC22-Spot-HA plasmid and selected by 25 µg/ml mycophenolic acid and 50 µg/ml xanthine.

### Construction of Apical-pellicle library

The gRNA sequences of Apical-pellicle library were selected from the genome-wide gRNA library^26^. The selected gRNA libraries were cloned into the modified pU6-Universal vector as previously described. The construction of the gRNA library was performed by VectorBuilder.

### *In vitro* and *in vivo* CRISPR screens

*In vitro* and *in vivo* CRISPR screens were conducted as previously described ^27^. Briefly, the gRNA library was linearized with NotI (Takara) and transfected into approximately 1–2×10^8^ RHΔhxgprt parasites divided between four separate cuvettes. Then, transfected parasites were grown in 4×150-mm dishes of confluent Vero cells. Medium was replaced with fresh one containing 10 µg/ml DNase I and 3 µM pyrimethamine 24 h post-transfection. Every 3 days until passage 3, parasites were scraped, mechanically lysed, and passaged to new dishes. After 2 days (passage 4), the parasites were syringe-lysed, filtered, and counted for genomic DNA preparation or for mouse infection. Parasites (1×10^8^) were pelleted and stored at-80 °C for genomic DNA preparation. The parasites were resuspended in PBS at a concentration of 2.5×10^8^ parasites/ml for mouse infection. Then, 1×10^7^ parasites in 40 µl PBS were injected into the footpad of each anesthetized mouse. Seven days post infection, mice were euthanized. Spleens were collected, homogenized by a plunger, and passed through a cell strainer to make cell suspensions. Then, the suspensions were added to 2×150-mm dishes per spleen with confluent Vero cells. The parasites typically lysed out after 2–4 days. At least 1×10^8^ parasites were pelleted and stored at-80 °C. Parasite genomic DNA was extracted using the DNeasy Blood and Tissue kit (Qiagen). The gRNAs were amplified and barcoded with KOD FX Neo (TOYOBO) using Primer 1 and Primer 2. The resulting libraries were sequenced on a DNBSEQ-G400RS (MGI) using Primer 3 and Primer 4. The primers used are shown in **Table S3**.

### Data analysis of the CRISPR screen

Following demultiplexing, gRNA sequencing reads were aligned to the gRNA library. The abundance of each gRNA was calculated and normalized to the total number of aligned reads. For *in vitro* analysis, the log_2_ fold-change between the passage 4 sample and the input library was calculated for each gRNA. For *in vivo* analysis, the log_2_ fold-change between each *in vivo* sample and the passage 4 sample was calculated for each gRNA. The median fitness score across mouse replicates was used as the *in vivo* fitness score. The p values were calculated using the Wilcoxon rank sum test between passage 4 vs *in vivo* sample. The adjusted p values were calculated using the Benjamini-Hochberg method. The distance of each gene from the regression line was calculated as below.

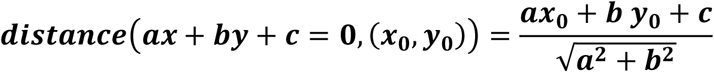

All analyses were conducted by R (v4.1.1) with package stats (v3.6.2) and visualized by ggplot2 (v3.4.0).

### Assessment of *in vivo* virulence in mice

Mice were infected with 10^3^ tachyzoites in 200 µl (intra-peritoneum) or 40 µl (intra-footpad) of PBS. The mouse health condition and survival were monitored daily until 30 days post infection.

### Immunofluorescence assay

HFFs were grown on coverslips, infected with parasites for 24–30 h, and fixed in 3.7% paraformaldehyde (PFA) for 10 min at room temperature. Cells were permeabilized with PBS containing 0.1% Triton-X for 10 min and then blocked with PBS containing 3% BSA for 1 h at room temperature. Then, the coverslips were incubated with the primary antibodies for 1 h, followed by incubation with appropriate secondary antibodies and DAPI for 30 min. The coverslips were mounted using PermaFluor (Thermo Scientific). Images were acquired by confocal laser microscopy (Olympus FV3000 IX83).

### Western blotting

Parasites were lysed in lysis buffer (1% NP-40, 150 mM NaCl, 20 mM Tris-HCl, pH 7.5) containing a protease inhibitor cocktail (Nacalai Tesque). The parasite lysates were separated by SDS-PAGE and transferred to polyvinylidene difluoride membranes (Immobilon-P; Millipore). Membranes were blocked in 5% skim milk in 0.05% Tween-20 in PBS for 1 h at room temperature, followed by incubation with primary antibodies for 1 h at room temperature. Membranes were incubated with appropriate secondary antibodies for 1 h at room temperature.

### Quantitative RT-PCR

Total RNA was extracted by RNeasy kit (Qiagen), and cDNA was synthesized by Verso reverse transcription (Thermo Fisher Scientific). Quantitative RT-PCR was performed with CFX Connect real-time PCR system (Bio-Rad) and Go-Taq real-time PCR system (Promega). The data was analyzed by the ΔΔCT method and normalized to ACT1 in each sample. The primers used are listed in **Table S3**.

### Microneme secretion assay

Freshly egressed parasites were pelleted and resuspended in serum-free DMEM with 2% ethanol for 30 min at 37°C. To suppress microneme secretion, parasites were also treated with 100 µM BAPTA-AM. After incubation, parasites were pelleted, and the supernatants were transferred to new collection tubes and re-centrifuged. The final supernatants (ESA) were resuspended in SDS loading buffer. The pellet lysates and ESA were subjected to immunoblot.

### Plaque assay

A total of 400 parasites were infected into a well of a six-well plate of HFFs and grown for 7 days in DMEM supplemented with 5% heat-inactivated FBS. Parasites were fixed with 3.7% PFA, stained with crystal violet, washed, and dried overnight. Plaque areas were measured by ImageJ.

### Invasion assay

Parasites were inoculated on coverslips with HFF monolayer in a 24-well plate. The plate was centrifuged for 5 min at 250 G, incubated for 1 h at 37°C, and fixed with 3.7% PFA. Extracellular parasites were stained with anti-SAG1 antibody without permeabilization. After washing three times, the cells were permeabilized with 0.1% Triton-X, and all parasites were stained with anti-GAP45 antibody. At least 100 parasites were counted and determined as extracellular or intracellular parasites.

### Replication assay

A total of 4×10^4^ parasites were inoculated on coverslips with HFF monolayer in a 24-well plate. Twenty-four hours post-infection, the monolayers were fixed with 3.7% PFA and processed for IFA with anti-GAP45 antibody to stain parasites. The number of parasites per vacuole was counted for at least 100 vacuoles.

### Induced egress assay

Freshly egressed parasites were inoculated on coverslips with HFF monolayer and grown for 24–30 hours at 37°C. For ionophore-induced egress assay, after washing with serum-free DMEM twice, the monolayers were incubated with 2 µM ionomycin (Nacalai Tesque) in serum-free DMEM at 37°C for 5 min. For low-pH induced egress assay, coverslips were washed twice with intracellular (IC) buffer (5 mM NaCl, 142 mM KCl, 1 mM MgCl_2_, 2 mM EGTA, 5.6 mM glucose, 25 mM HEPES, pH adjusted to 7.4 with KOH) and the buffer was replaced with IC buffer (pH adjusted to 5.4 or 7.4 with HCl) with or without 0.002% digitonin, and coverslips were incubated at 37°C for 3 min. Cells were fixed with 3.7% PFA and processed for IFA with anti-SAG1 and anti-GRA17 antibodies to stain parasites and PVs, respectively. At least 100 vacuoles were counted per strain and scored as occupied or egressed.

### Gliding motility assay

Freshly egressed parasites were resuspended in serum-free DMEM and added to poly-L-Lysine coated coverslips in a 24-well plate. The plate was centrifuged for 3 min at 250 G. The media was replaced with serum-free media containing 2% EtOH. Following incubation for 30 min at 37°C, the coverslips are fixed with 3.7% PFA and processed for IFA with anti-SAG1 antibody to stain SAG1 trails.

### Time-lapse imaging of membrane permeabilization

Parasites were inoculated on HFF monolayers in glass-bottom culture dishes and grown for 24–30 hours at 37°C. HFF monolayers were pre-treated with 2 µM Cytochalasin D in DMEM for 5 min followed by the addition of an equal volume of 2 µM Cytochalasin D and 4µM ionomycin in DMEM.

Time-lapse images were acquired every 10 sec for up to 10 min by confocal laser microscopy (Olympus FV1200).

### Detection of PLP1 dynamics induced by ionophore treatment

A total of 2×10^6^ parasites were inoculated on monolayers of human fibroblasts in six-well plates and grown for 24 h at 37°C. The monolayers were pre-treated with 2 µM Cytochalasin D in DMEM for 5 min followed by the addition of an equal volume of 2 µM Cytochalasin D and 4µM ionomycin in DMEM for additional 5 min. Parasites and host cells were lysed in lysis buffer and subjected to immunoblots.

## Co-immunoprecipitation followed by mass spectrometry

ΔMIC11::MIC11-Spot-HA parasites or the parental ΔMIC11 parasites were fixed with 0.1% PFA and lysed in HEPES-RIPA buffer (20 mM HEPES-NaOH, pH 7.5, 1 mM EGTA, 1 mM MgCl_2_, 150 mM NaCl, 0.25% sodium deoxycholate, 0.05% SDS, 1% NP-40) supplemented with protease inhibitor cocktail cOmplete EDTA-free (Roche) and Benzonase (Merck). After centrifugation, the supernatants were incubated with Spot-Trap magnetic agarose beads (Proteintech, etma) for 3 h at 4°C. The beads were washed three times with HEPES-RIPA buffer and twice with 50 mM ammonium bicarbonate. Proteins on the beads were digested with 200 ng of trypsin/Lys-C mix (Promega) at 37°C overnight. The digested peptides were reduced, alkylated, acidified, and desalted using GL-Tip SDB (GL Sciences). The eluates were evaporated in a SpeedVac concentrator and dissolved in 0.1% trifluoroacetic acid and 3% acetonitrile (ACN). LC-MS/MS analysis of the resultant peptides was performed on an EASY-nLC 1200 UHPLC connected to an Orbitrap Fusion mass spectrometer through a nanoelectrospray ion source (Thermo Fisher Scientific). The peptides were separated on a 75 µm inner diameter × 150 mm C18 reversed-phase column (Nikkyo Technos) with a linear 4–32% ACN gradient for 0–100 min followed by an increase to 80% ACN for 10 min and a final hold at 80% ACN for 10 min. The mass spectrometer was operated in a data-dependent acquisition mode with a maximum duty cycle of 3 s. MS1 spectra were measured with a resolution of 120,000, an automatic gain control (AGC) target of 4e5, and a mass range from 375 to 1,500 m/z. HCD MS/MS spectra were acquired in the linear ion trap with an AGC target of 1e4, an isolation window of 1.6 m/z, a maximum injection time of 35 ms, and a normalized collision energy of 30. Dynamic exclusion was set to 20 s. Raw data were directly analyzed against *T. gondii* GT1 protein data (ToxoDB release 55) supplemented using Proteome Discoverer version 2.5 (Thermo Fisher Scientific, USA) with Sequest HT search engine. The search parameters were as follows: (a) trypsin as an enzyme with up to two missed cleavages; (b) precursor mass tolerance of 10 ppm; (c) fragment mass tolerance of 0.6 Da; (d) carbamidomethylation of cysteine as a fixed modification; and (e) acetylation of protein N-terminus and oxidation of methionine as variable modifications. Peptides and proteins were filtered at a false discovery rate (FDR) of 1% using the percolator node and the protein FDR validator node, respectively. Label-free precursor ion quantification was performed using the precursor ions quantifier node, and normalization was performed such that the total sum of abundance values for each sample over all peptides was the same.

### Production of recombinant proteins

The N-terminal His-tagged recombinant MIC11-HA and PLP1-FLAG were expressed using the wheat germ cell-free expression system (CellFree Sciences, Matsuyama, Japan), and purified by immobilized-nickel affinity chromatography with Ni Sepharose™ 6 Fast Flow (Cat#17531802, Cytiva).

### Co-immunoprecipitation of recombinant proteins

Recombinant proteins (0.5 µg) were co-incubated in equilibrium buffer (20 mM Tris-HCl, pH 7.5, 250 mM NaCl) at 4°C for 1 h. Anti-HA antibody-agarose was added, and proteins were further incubated at 4°C for 1 h. The protein solution was centrifuged and washed with Tris-buffered saline (TBS) three times. Immunoprecipitated proteins were eluted by boiling with SDS loading buffer and subjected to immunoblot.

### Homologue search and sequential alignment

VEuPathDB^69^ was searched for MIC11 homologues. MAFFT^70^ was used to align and visualize MIC11 and MIC22.

### Quantification and statistical analysis

All statistical analyses except for the survivals were performed using R (4.1.1, https://www.r-project.org/). Pearson’s correlation was used for correlation analysis. Data with p values < 0.05 were considered statistically significant. The statistical analysis of survival rates was performed by the log-rank test using the GraphPad Prism9 software. All images shown are representative results from one of two or more experiments.

## Data availability

The CRISPR screen data has been deposited to the NCBI GEO under the accession number **GSE290078**. Any additional information required to reanalyze the data reported in this paper is available from the corresponding author upon request.

## Acknowledgements

We thank M. Enomoto and N. Yamagishi (Osaka University) for the secretarial and technical assistance. We acknowledge the NGS core facility at the Research Institute for Microbial Diseases of Osaka University for the sequencing. This study was supported by Japan Science and Technology Agency (JPMJFR206D and JPMJMS2025); Agency for Medical Research and Development (JP25wm0325072, JP223fa627002, and JP25fk0108682); Ministry of Education, Culture, Sports, Science and Technology (23KJ1469); the program from Joint Usage and Joint Research Programs of the Institute of Advanced Medical Sciences, Tokushima University; Takeda Science Foundation; Mochida Memorial Foundation; Astellas Foundation for Research on Metabolic Disorders; Naito Foundation; the Chemo-Sero-Therapeutic Research Institute; Research Foundation for Microbial Diseases of Osaka University; BIKEN Taniguchi Scholarship; The Nippon Foundation - Osaka University Project for Infectious Disease Prevention; Joint Research Program of Research Center for Global and Local Infectious Diseases of Oita University (2021B06).

## Author contributions

Conceptualization, Y.T. and M.Y.; Methodology, Y.T. and M.Y.; Software and formal analysis, Y.T.; Investigation, Y.T., H.K., and M.Y.; Writing – Original Draft, Y.T. and M.Y.; Writing – Review & Editing, Y.T., M.S., V.B.C., D.S.F., and M.Y.; Funding Acquisition, Y.T. and M.Y.; Resources, E.T., V.B.C., and D.S.F.; Supervision, V.B.C., D.S.F., and M.Y.

---

**Table S1**

Summary of CRISPR screen, raw count data, fitness scores, and gRNA sequences for each gene.

**Table S2**

Summary of IP-MS analysis.

**Table S3**

List of primers used in this study.

## Extended figures

**Figure S1.**
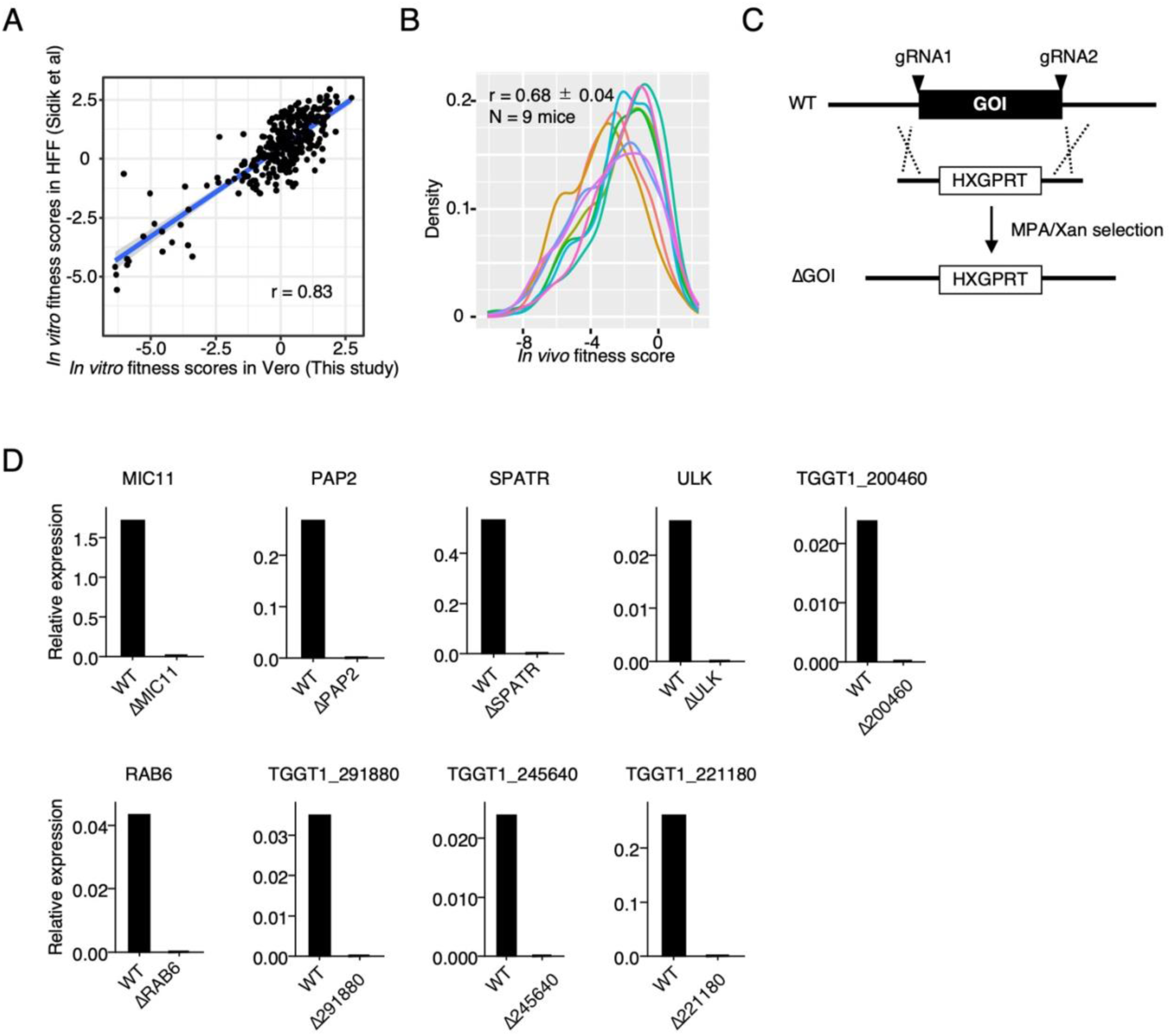
Validation of the screen and knockout parasites, related to. Figure 1. (A) Correlation between the *in vitro* fitness scores in Vero and the *in vitro* fitness scores in HFF (Sidik et al., 2016). (B) Overlay of *in vivo* fitness scores for each mouse. Pearson’s correlation coefficients are shown as mean ± SD. (C) Schematic of gene knockout strategy. (D) Quantitative RT-qPCR validations for indicated knockout parasites.

**Figure S2.**
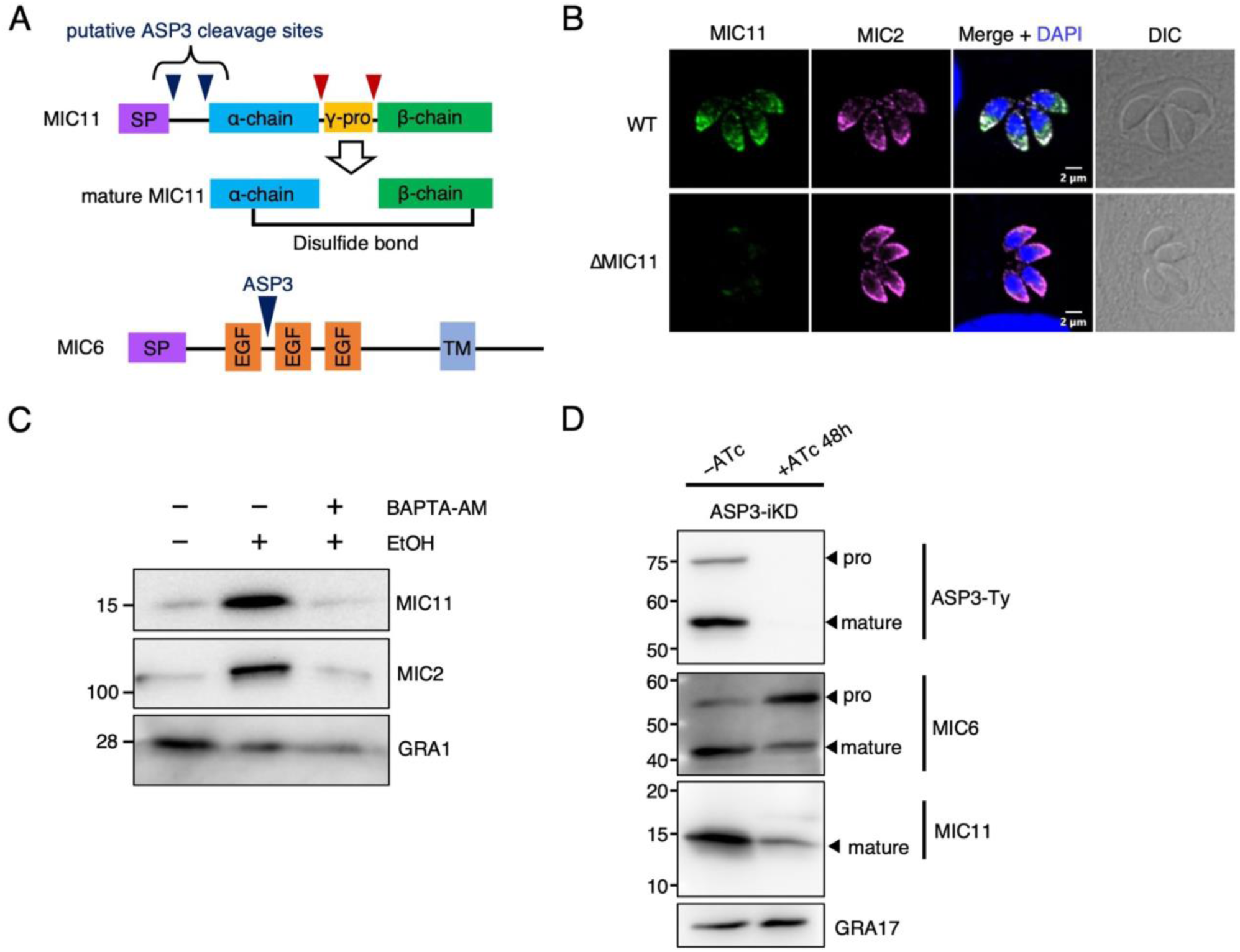
MIC11 is a microneme protein processed by ASP3. (A) Schematic representation of MIC11 and MIC6. Their putative cleavage sites by ASP3 were shown in dark blue. (B) Immunofluorescence images stained with anti-MIC11 (green) and anti-MIC2 (magenta) antibodies. (C) Microneme secretion assay showed MIC11 was secreted in a Ca^2+^-dependent manner. (D) Immunoblots assessing the processing of microneme proteins, MIC6 (a known ASP3 substrate) and MIC11, upon ASP3 inducible knockdown. GRA17 was used as the loading control.

**Figure S3.**
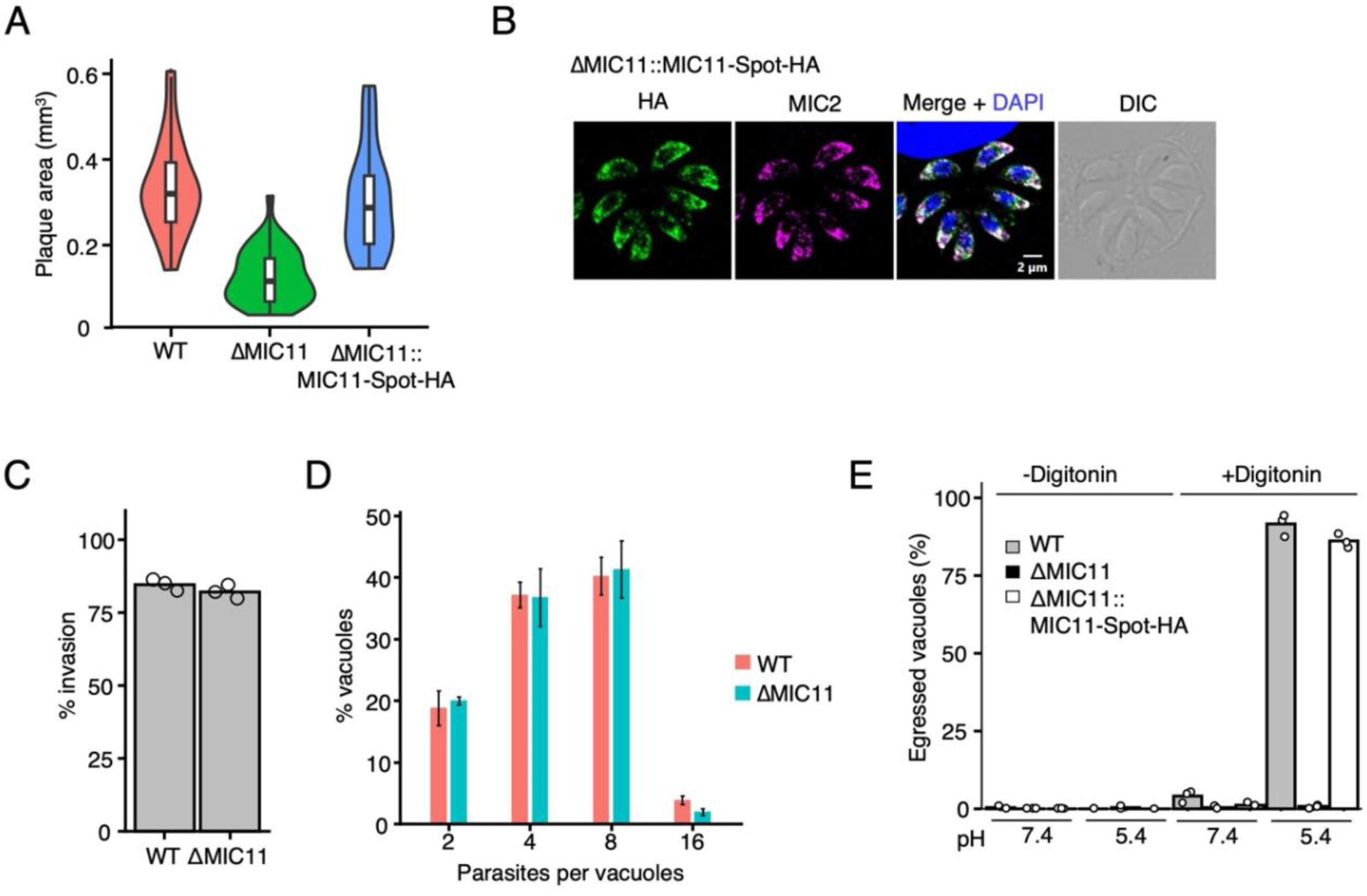
Phenotypic analysis of ΔMIC11 parasites, related to Figure3. (A) Plaque areas of indicated strains. (B) Immunofluorescence images of MIC11::MIC11-Spot-HA parasites. (C) Invasion assay. (D) Intracellular replication assay. (E) Low pH-induced egress assay of indicated strains.

**Figure S4.**
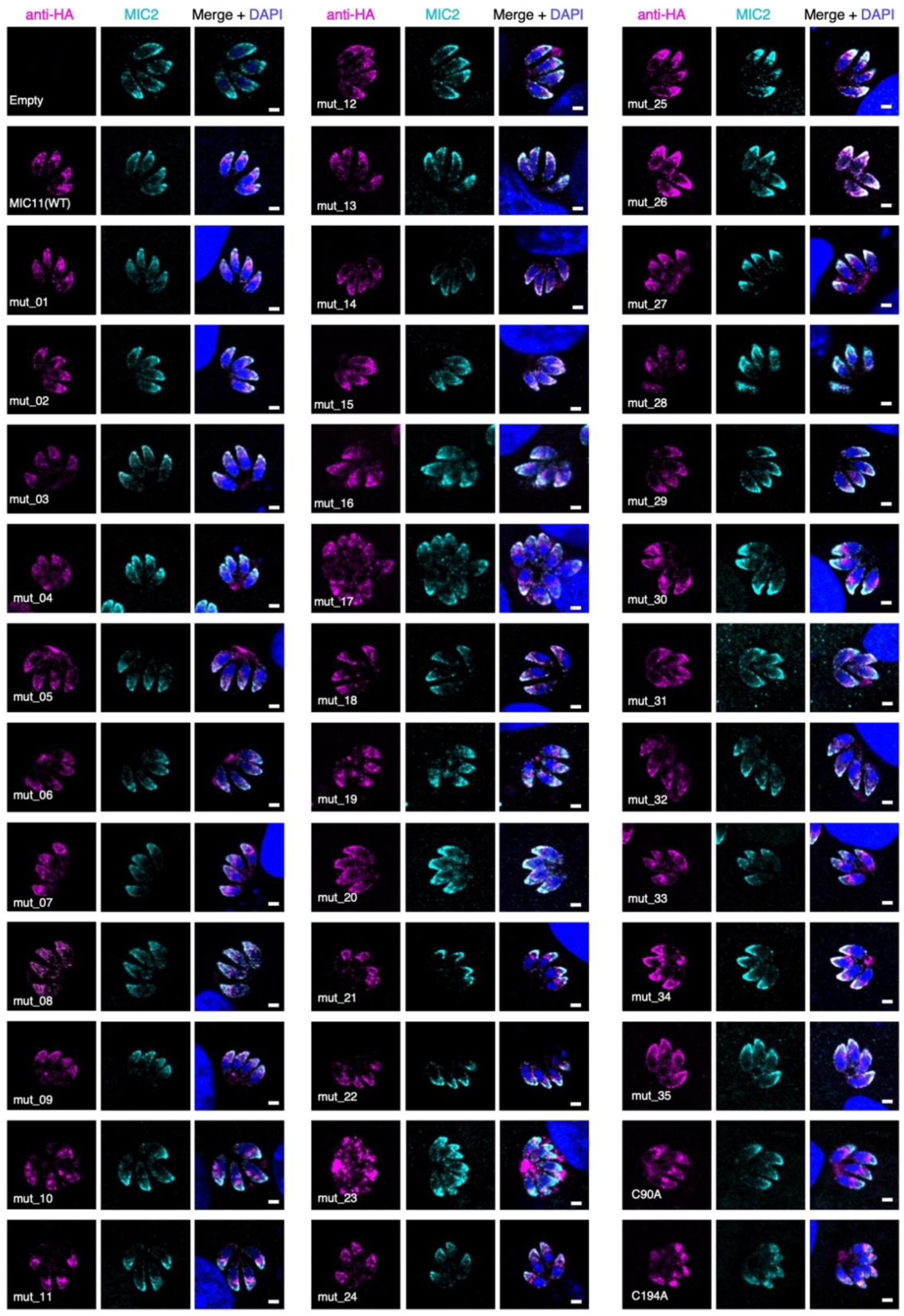
Validation of MIC11 localization, related to Figure4. Scale bar = 2 µm for all.

**Figure S5.**
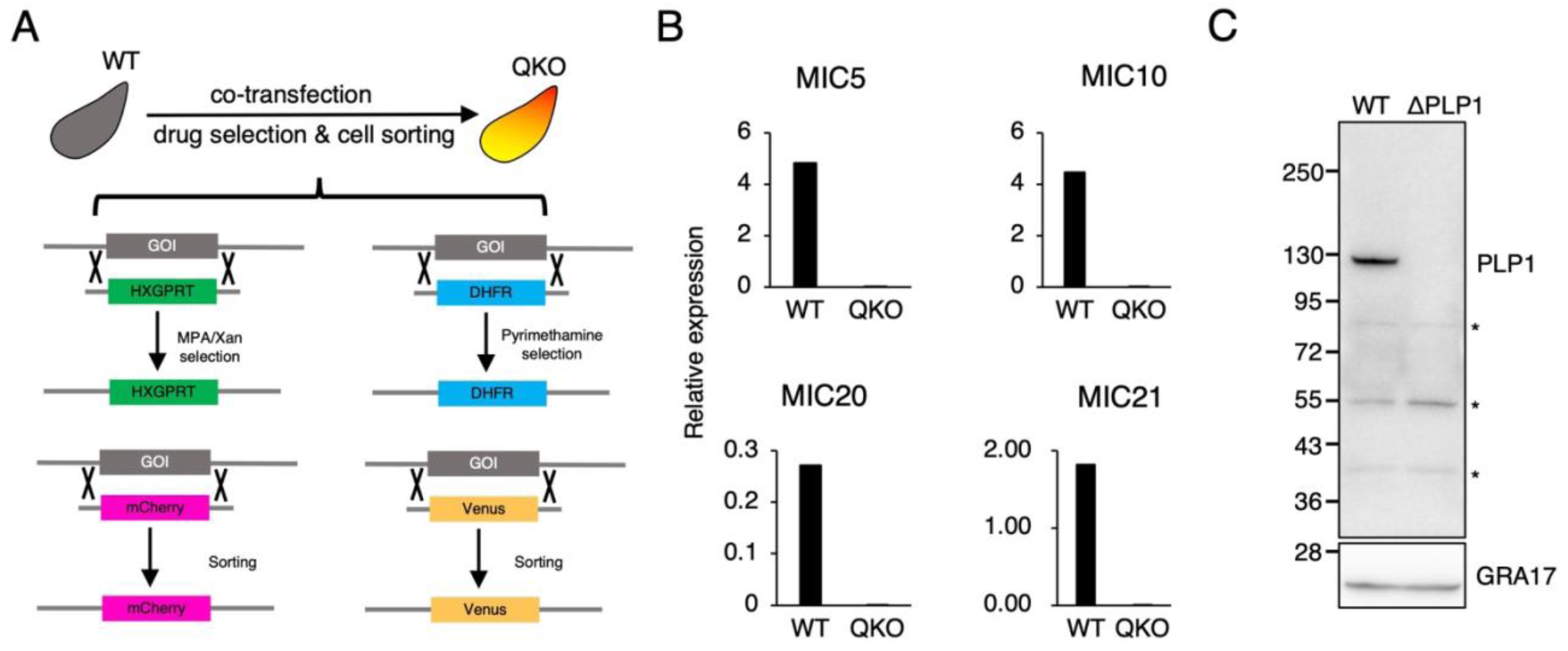
Generation of ΔPLP1 and QKO parasites, related to. Figure 5. (A) Schematic of quadruple knockout. (B) Validation of quadruple knockout by RT-qPCR. (C) Validation of PLP1 knockout by immunoblot. Asterisks indicate non-specific bands.

**Figure S6.**
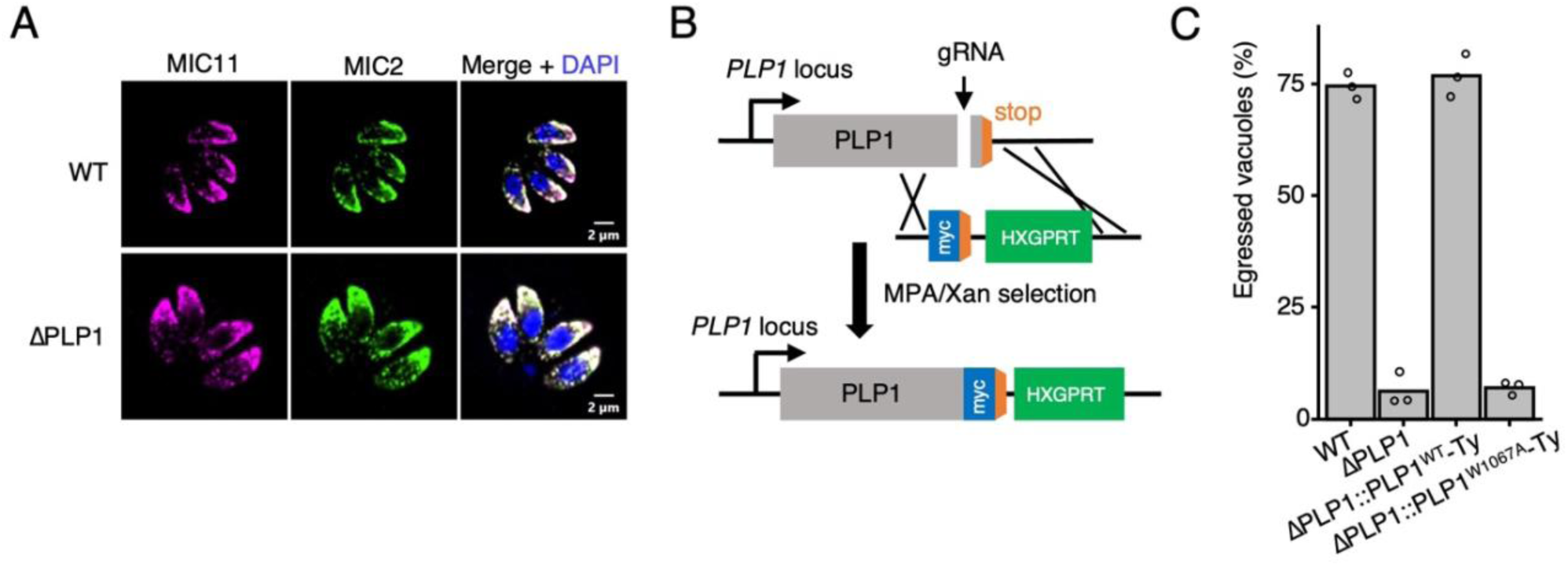
MIC11 localization is not affected by PLP1 deletion, related to. Figure 6. (A) Immunofluorescence images stained with anti-MIC11 (magenta) and anti-MIC2 (green) antibodies. (B) Schematic of PLP1 endogenous tagging. (C) Induced-egress assay of indicated strains.

